# Likelihood analysis of population genetic data under coalescent models: computational and inferential aspects

**DOI:** 10.1101/185702

**Authors:** François Rousset, Champak Reddy Beeravolu, Raphaël Leblois

## Abstract

Likelihood methods are being developed for inference of migration rates and past demographic changes from population genetic data. We survey an approach for such inference using sequential importance sampling techniques derived from coalescent and diffusion theory. The consistent application and assessment of this approach has required the re-implementation of methods often considered in the context of computer experiments methods, in particular of Kriging which is used as a smoothing technique to infer a likelihood surface from likelihoods estimated in various parameter points, as well as reconsideration of methods for sampling the parameter space appropriately for such inference. We illustrate the performance and application of the whole tool chain on simulated and actual data, and highlight desirable developments in terms of data types and biological scenarios.

**Résumé:** Diverses approches ont été développées pour l’inférence des taux de migration et des changements démographiques passés à partir de la variation génétique des populations. Nous décrivons une de ces approches utilisant des techniques d’échantillonnage pondéré séquentiel, fondées sur la modélisation par approches de coalescence et de diffusion de l’évolution de ces polymorphismes. L’application et l’évaluation systématique de cette approche ont requis la ré-implémentation de méthodes souvent considérées pour l’analyse de fonctions simulées, en particulier le krigeage, ici utilisé pour inférer une surface de vraisemblance à partir de vraisemblances estimées en différents points de l’espace des paramètres, ainsi que des techniques d’échantillonage de ces points. Nous illustrons la performance et l’application de cette série de méthodes sur données simulées et réelles, et indiquons les améliorations souhaitables en termes de types de données et de scénarios biologiques.

**Mots-clés:** histoire démographique, processus de coalescence, importance sampling, genetic polymorphism

**AMS 2000 subject classifications:** 92D10, 62M05, 65C05

## 1. Introduction

Since the advent of genetic markers, there have been many efforts to infer demographic parameters (population sizes and dispersal) from observed genetic variation. These efforts serve to better understand the forces affecting the evolution of natural population, and also appear to fulfill a distinct fascination for the history of past human migrations and population admixtures. Early statistical approaches have considered descriptions of genetic variation that can be understood as analyses of variance in allele frequencies among different groups of individuals (Cockerham, 1973). In particular, Wright’s *F*-statistics (Wright, 1951) can be expressed as functions of frequencies of pairs of gene copies that are of identical allelic state, and then viewed as estimators of the corresponding functions of probabilities that pairs of gene copies are identical, under a given model. As there are theoretical expectations for these probabilities in simple models of evolution, a quantitative process-based interpretation of the descriptors is possible, to infer dispersal parameters among different subpopulations, or the demographic history of natural populations.

For the same objectives, likelihood analyses attempt to extract information from the joint allelic types of more than two genes copies. These attempts have been hampered by the increasing difficulty in computing the probability distribution of such joint configurations as the number of gene copies increases. For this reason, stochastic algorithms have been developed to estimate the likelihood of a sample of arbitrary size. These algorithms view a sample as incomplete data, where the missing information is the genealogy of all gene copies in the sample. If the “complete-data” likelihood, that is the probability of the sample given the genealogy, is easy to evaluate, the evaluation of the sample likelihood can be formulated as the evaluation of a marginal likelihood, obtained by integration of this complete-data likelihood over a probability distribution of genealogies consistent with the data. A classic recurrent Markov chain Monte Carlo approach has been used to sample from this distribution (Beerli and Felsenstein, 1999; Nielsen and Wakeley, 2001; Hey, 2010). However, the slow convergence of such methods has prompted both the development of alternative algorithms for computing the marginal likelihood, and also explains the persistence of the older methods and the development of other methodologies based on simulation of samples, such as Approximate Bayesian Computation (Beaumont, 2010).

In this paper we review an approach to perform likelihood-based inferences, using a class of importance sampling algorithms derived from the work of Griffiths and collaborators (de Iorio and Griffiths, 2004a,b; see also Stephens and Donnelly, 2000). We first explain the importance sampling algorithm defined in this work to obtain estimates of the likelihood of given parameter points. Next we discuss the additional steps required to derive reliable inferences from such likelihood estimates. A distinctive feature of the latter work, when compared to most of the literature on alternative methods of inference, is the emphasis on evaluating the inference in terms of coverage properties of likelihood-based confidence intervals. For such purposes, one has to infer a likelihood surface from estimated likelihoods in different parameter points. Kriging has classically been used for inference of response surfaces (e.g., Sacks et al., 1989), and our efforts to obtain good coverage has led us to reimplement such methods as part of a set of software tools to explore likelihood surface in an automatic way.

The methods described in this paper are all implemented in free software: the MIGRAINE software, written in C++, implements the algorithms for likelihood estimation in each given parameter point, and calls R code that performs inference of the likelihood surface from the likelihood points, plots various representations of this surface and other diagnostics, evaluates likelihood ratio confidence intervals, and designs new parameter points which likelihood should be computed in a next iteration of MIGRAINE. Most of this R code has recently been incorporated in a standard R package, blackbox, which can also be used on its own to perform optimization of simulated functions. MIGRAINE writes all the required calls to R functions so that no understanding of them is required from the user.

## 2. Lihelihood inference using importance sampling algorithms

### 2.1. Inferring the likelihood for a parameter point by importance sampling

#### 2.1.1. Sequential importance sampling formulation

Given a current sample **S**, we consider the ancestral states (i.e. allelic types) of the gene lineages ancestral to **S** at any time *t*, called the “ancestral sample”, **S**(*t*). These ancestral states are considered at any time until the time *t*_*τ*_ where a common ancestor of the sample is reached. We consider transition probabilities 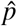 for **S**(*t*_*k*_) over successive time steps *t*_0_*,t*_1_*, …, t*_*τ*_, and importance sampling weights 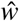 defined such that the likelihood of a sample can be written as

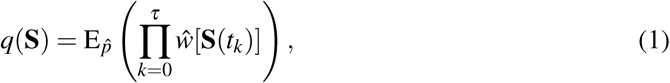

where the expectation is taken over the distribution of sequences (**S**(*t*_*k*_)) generated by the transition probabilities 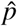. These transition probabilities define a Markov chain over ancestral states, with absorbing states being reached at time *t*_*τ*_ when a single common ancestor is reached. Estimation of *q*(**S**) is then performed by averaging 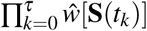 over independent realizations of this Markov chain (typically 2000 such independent genealogies in the following applications).

de Iorio and Griffiths (2004a,b) propose 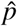 and 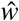 based on approximations for the ratio *π ≡ q*(**S**)*/q*(**S**′) of the probabilities of samples differing by one event (mutation, migration, or coalescence event). We will detail how these approximations are constructed. For that purpose we will first consider recursions over a time interval, relating the current sample to an ancestral sample **S**(*t*) taken (say) a generation before.

These recursions are obtained by a coalescent argument. That is, we represent the events leading to the current sample of *n* genes as the realizations of two processes: a coalescent process determining the marginal distribution of ancestral genealogies of *n* genes, independent of the current allelic types; and given a genealogy, a mutation process that changes the allelic types along the branches of the genealogical tree. For developments of coalescent methods see Tavaré (1984), Hein et al. (2005), or Wakeley (2008).

In this perspective, the relationship between a current sample probability and the parental sample probability can be conceived as the joint realizations of two processes in addition to those leading to the parental sample: the marginal genealogical process over the latest generation, and the mutation process over this generation. In the following we consider samples from subdivided populations, where sample size is defined as a vector **n** of sample sizes in distinct subpopulations, and samples are characterized by the counts of different alleles in each sampled subpopulations. The recursion between a current sample **S**′ and all possible parental samples **S** takes the form

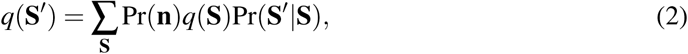

where *q*(**S**′) Pr(**S**′ **n**′) is the stationary probability that the descendant sample is **S**^*t*^, given the descendant sample size **n**′; *q*(**S**) Pr(**S n**) is likewise the stationary probability of sample **S** given parental sample size **n**; Pr(**n**) Pr(**n n**′) is the stationary probability that, given the descendant **n**′ (but not given **S**′, by the coalescent argument), the parental lineages form a sample of **n** genes. It is thus the stationary probability of genealogical events in the latest generation; and Pr(**S**′ **S**) Pr(**S**′ **S**, **n**′) is the probability (given **n**′) tht mutation events led to the descendant sample **S**′ given the parental sample **S** and the descendant **n**′.

This recursion suggests the following inefficient importance sampling algorithm. We rewrite the recursion by discarding the case where **S**′ = **S** on the right-hand sum. The resulting equation can be written as

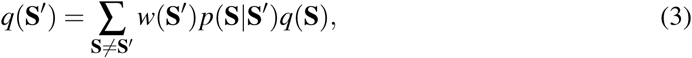

where

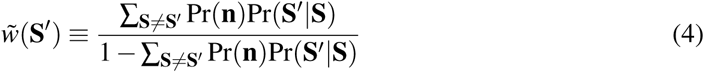

and

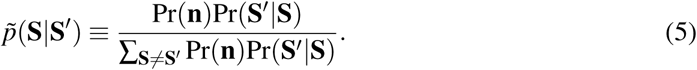

The probabilities 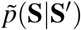 define transition probabilities of a Markov chain such that

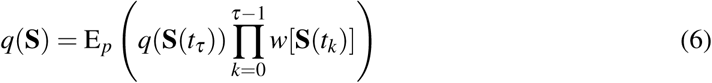

where **S**(*t*_0_) = **S** represents the allelic counts in the current sample, and **S**(*t*_*τ*_) the allelic type of the most recent common ancestor of **S**(*t*_0_). Thus, the 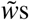 (or their product) are importance sampling weights in a sequential importance sampling algorithm of which the proposal distribution is the distribution of ancestral histories generated by 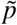.

A good pair (*p, w*) is such that 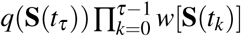 has low variance over realizations of *p*. The above pair is inefficient in this respect. An optimal IS algorithm can be defined as yielding a zero variance, and Stephens and Donnelly (2000) characterized the optimal pair (*p, w*) in terms of successive samples and their stationary probabilities. To derive a feasible algorithm from this characterization, de Iorio and Griffiths (2004a,b) reformulated it in terms of the probabilities *π*(*j d,* **S**), for any *j* and *d*, that an additional gene taken from subpopulation *d* is of type *j*. Then, approximations for the optimal (*p, w*) can be defined from approximations for the *π*s.

#### 2.1.2. Optimal p and w

Rewrite

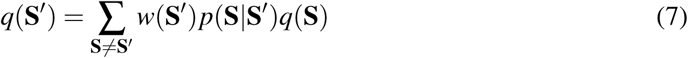

as

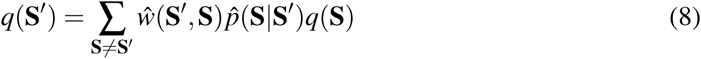

for some transition probabilities 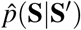 forming a Markov transition matrix, and for

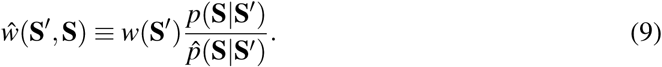

Then 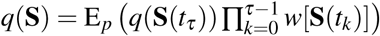 becomes 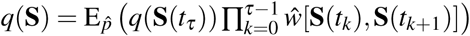

Consider the Markov chain defined by the transition probabilities

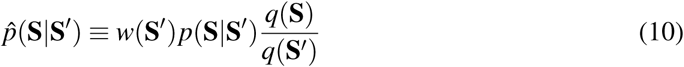

for any pair **S**′, **S**. Then 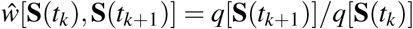 and any realization of this Markov chain over ancestral states gives the exact likelihood (“perfect simulation”):

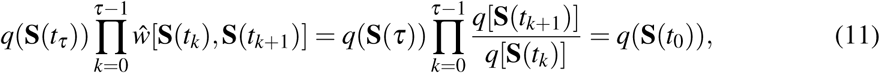

which shows that 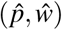 is optimal.

#### 2.1.3. Formulation of efficient p and w

We can rewrite the optimal importance sampling algorithm in terms of the probability *π*(*j |d,* **S**) that an additional gene taken from deme *d* is of type *j* (such that the sum over all possible types ∑ _*j*_ π(*j |d,* **S**) = 1). We write the stationary probability *q*(**S**) as an expectation over the joint distribution of frequencies *X*_*di*_ for all alleles *i* in all subpopulations *d*,

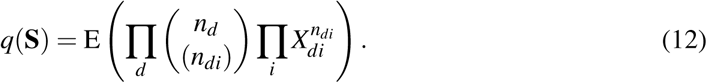

Then for any *d* and *j*, *π*(*j|d,* **S**) is related to the stationary sample probabilities by

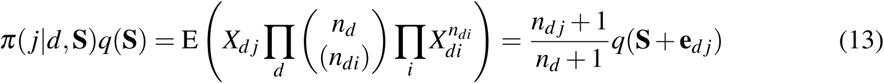

where the expectation is taken over the stationary density of joint allele frequencies **x** in the different demes considered. Thus if two successive samples differ by the addition of a gene copy of type *j* in deme *d*, the corresponding term 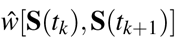corresponding term in eq. 11 can be written as

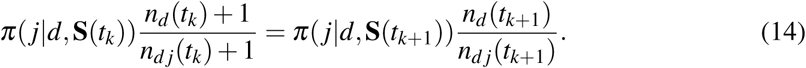

If two successive samples differ by a mutation from *i* to *j* in deme *d*, then

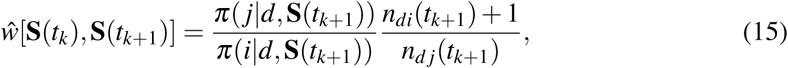

as mutation can be represented as the removal of one gene copy and the addition of another gene copy of another type in the same deme. Likewise, a migration from deme *d* to deme *d*^*t*^ yields

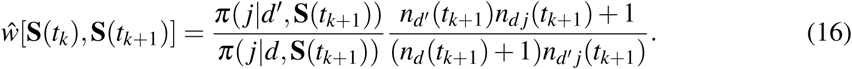

Coalescent methods typically consider that only one event (coalescence, mutation, or migration) distinguishes the successive samples. Thus, in informal terms, the mutation and migration rates are assumed small, and subpopulation sizes are assumed large, so that it is unlikely that more than one coalescence event occurs in a generation (see the Appendix for a somewhat more formal statement). Then, the product of sequential weights in eq. 11 can be written, for any sequence of ancestral samples, as a product of terms given in the last three equations. Any approximation for the *π*s then defines an approximation for the optimal weights in an importance sampling algorithm.

The Appendix details the approximation defined by de Iorio and Griffiths (2004a,b). This approximation recovers the true *π*s and thus allows “perfect simulation” in a few cases where the stationary distribution of allele frequencies in populations is known, and it is otherwise very efficient for other time-homogeneous models that have been investigated (de Iorio et al., 2005; Rousset and Leblois, 2007, 2012). The previous arguments also yield importance sampling algorithms for time-inhomogeneous models where the rates of events depend on time-variations in parameter values, when random times are attached to the successive events in the ancestral history (Griffiths and Tavaré, 1994). The 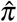 approximation of de Iorio and Griffiths (2004a,b) has been used to extend the inference method to models with changing population size over time (Leblois et al., 2014) and models with population divergence events (divergence with migration between two populations, unpublished work). However, the 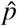 proposal at any step *t* only takes into account the rates at time *t*, not the more ancestral rate variations that also affect sample probabilities at time *t*, and this results in a loss of efficiency of the IS algorithm. Resampling methods (Liu, 2004) have been investigated to provide some relief to this inefficiency (Merle et al., 2017).

#### 2.1.4. The PAC-likelihood heuristics

Eq. 11 holds for any sequence (**S**(*t*_*k*_)), even if this sequence is not a biologically coherent sequence of ancestral states. Thus it holds for any sequence *S*_*l*_ defined as the sequential addition of all constituent gene copies *g*_*l*_ (*l* = 1*, …, n*) of the final sample **S**, in any order. For such a sequence eq. 11 takes the form

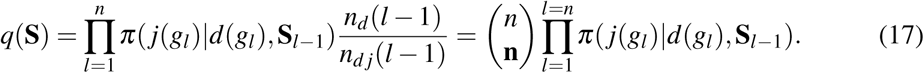

where *j*(*g*_*l*_) and *d*(*g*_*l*_) represent respectively the allelic type of gene copy *g*_*l*_ and the subpopulation where it is added. Using some approximation for the *π*s in this expression yields a Product of Approximate Conditional (PAC) approximation to the likelihood (Li and Stephens, 2003). It is heuristic, in the sense that it is generally not a consistent estimator of the likelihood. However, we can use the same approximations to the *π*s as in the importance sampling algorithm (Cornuet and Beaumont, 2007), and in that case likelihood inference based on PAC-likelihood has proven practically equivalent to that based on likelihood (Rousset and Leblois, 2007, 2012; Leblois et al., 2014). The main drawback of the approximation is that, since there is no ancestral time attached to the successive **S**(*l*), this PAC-likelihood approximation cannot substitute the IS approach in models with time-varying rates. However, some models with time-varying rates include a ancestral stable demographic phase (e.g. the model with a contraction or an increase in population size used in Leblois et al. (2014) and illustrated in Fig. 1). Under such models, the PAC-likelihood can still be used to approximate the probability of the states of the ancestral genes lineages remaining when the stable demographic phase is reached backwards in time, and this approximation has allowed significant decreases in computation time without loss in precision (Leblois et al., 2014).

**Figure 1:**
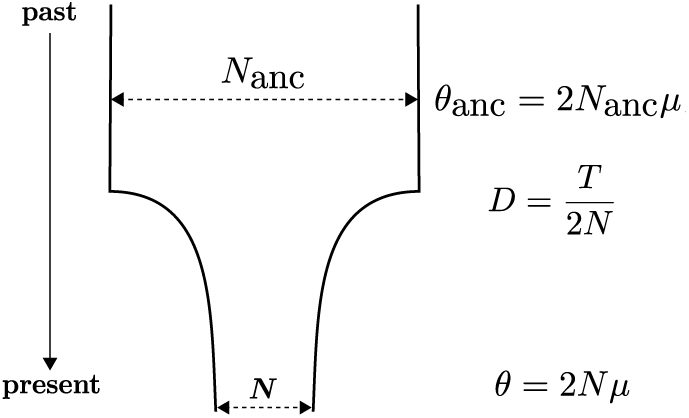
Representation of the time-inhomogeneous demographic model considered in Leblois et al. (2014). *N* is the current population size, *N*_anc_ is the ancestral population size (before the demographic change), *T* is the time measured in generations since present and *µ* the mutation rate of the marker used. Those four parameters are the parameters of the finite population model. *θ*, *D* and *θ*_anc_ are the inferred scaled parameters of the coalescent approximations.

### 2.2. Inferring the likelihood surface by smoothing

The above algorithms provide estimates of likelihood for given parameter values. A difficulty encountered in the first applications of this methodology (de Iorio et al., 2005) is that widely usable software (in particular, various R packages) were not up to the task of accurately inferring a likelihood surface from a collection of such estimates, and that the best of them still failed in a notable fraction of computations (essentially in the inversion of near-singular matrices), hampering our validation efforts. Our re-implementation of Kriging uses generalized cross-validation (Golub et al., 1979; Nychka, 2000) to obtain estimates of the smoothing parameters in reasonable time, in a way similar to the fields package in R (Nychka et al., 2015). It also uses a complex strategy (discussed below) to sample points in parameter space in an automated way with minimal input from users. The details of the sampling strategy can substantially impact the performance, particularly as the number of parameters increases, but this impact cannot be fully assessed unless performance of the overall inference method (e.g., coverage of confidence intervals in the present case) is itself assessed.

To obtain a first estimate of the likelihood surface, one has to sample evenly in parameter space. However, to estimate smoothing parameters, clusters of close parameters points are also useful (Zimmerman, 2006). The current implementation performs an empirical compromise between these distinct needs. From any estimate of the likelihood surface, further parameter points can then be sampled. The general resampling strategy, as detailed below, is to define a space of parameters with putatively high likelihood according to the current likelihood surface estimate, then to sample at random within this space, and to select among the sampled points those that are appropriate or best according to some additional criteria. MIGRAINE allows extrapolation beyond the parameter regions sampled in previous iterations, subject to ad hoc constraints in parameter space (such as positive mutation rates, but sometimes more complex constraints for composite parameters such as the so-called neighborhood size in models of localized dispersal).

Part of the new points are sampled uniformly in a parameter region with high predicted likelihood. But parameter regions that have yet been little sampled typically have high prediction variance, and may thus be worth sampling even if the predicted likelihood is relatively low in such regions. Expected improvement (EI) methods allow sampling of points in such regions by taking in account both point prediction and high variance in prediction (e.g., Bingham et al., 2014). The latest versions of MIGRAINE use EI to generate part of the new points, by first sampling a larger number of points (typically 100 times the target number) uniformly in a given parameter region, then retaining the ones with best EI. This approach is used to more accurately identify the ML estimates, but also the confidence limits. In the latter case, confidence limits (*λ_−_, λ*_+_) for any parameter *λ* are deduced from the profile log-likelihood ratio (LR) defined by maximization over other parameters *ψ*. Then EI is used to select new values of *ψ* given *λ* = *λ*_*−*_ or *λ* = *λ*_+_. Additional points with high EI are also selected specifically outside the parameter regions with highest predicted likelihood.

A nice feature of this iterative approach is that it is not very important to have accurate estimation of likelihood in each parameter point, because the accumulation of likelihood estimates nearby the maximum (or any other target point) over successive iterations will provide, by the infill asymptotic properties of Kriging (Stein, 1999), an accurate estimation of likelihood at the maximum. For example, under models of localized dispersal (so-called “isolation by distance” in population genetics), simulating 20 genealogies per parameter point is sufficient to obtain almost perfect coverage properties of the confidence intervals (Rousset and Leblois, 2012).

## 3. Examples

### 3.1. Inference of a founder event in Soay sheep

We will illustrate the whole inference procedure (i.e. likelihood computation at different points of the parameter space and likelihood surface smoothing) by analyzing available data from an isolated sheep population from the island of Hirta, previously published in Overall et al. (2005). The Hirta island was evacuated of humans and their modern domestic sheep in 1930, and 107 sheep were reintroduced in 1932 from the neighboring island of Soay. The population has since remained unmanaged and the total island population has been recently observed to reach up 2000 individuals. The data set consists in 198 individual genotypes, thus 396 gene copies, screened at 17 microsatellite markers. All genotyped individuals were born in 2007.

For this application, we first considered the model of a single population with a single past change in population (i.e. the model presented in Fig. 1), which has been thoroughly tested by simulation in Leblois et al. (2014) and applied on different data sets (e.g. Vignaud et al., 2014a; Lalis et al., 2016; Zenboudji et al., 2016). Microsatellite alleles are repeats of a very short DNA motif, and mutation models generally describe the distribution of change in number of repeats when a mutation occurs. The model assumed here is the generalized stepwise mutation model (GSM, Pritchard et al., 1999) characterized by a geometric distribution of mutation steps, with parameter *p*_GSM_. Four (scaled) parameters are thus inferred under this model: *p*_GSM_, *θ* = 2*Nµ*, *D* = *T /*2*N*, and *θ*_anc_ = 2*N*_anc_*µ* (see legend of Fig. 1 for explanation of the model parameters). An additional composite parameter, *N*_act/anc_ = *N/N*_anc_, describes past changes in population size (i.e. past contraction or expansion). The present analysis consisted in 8 iterations, each with 200 parameter points. For each point the likelihood is estimated using 2,000 genealogies. Initial parameter ranges as well as point estimates and associated confidence intervals (CI) are presented in Table 1 (lines ‘Single change‘) and examples of one- and two-dimensional profile likelihood ratio (LR) plots are shown in Fig. 2. In all such plots, the likelihood profile are inferred only from the likelihood of parameters restricted within the convex hull of sampled parameter points, i.e. ignoring values inferred by the Kriging prediction outside this region. There is enough information on all parameters in the data set for the analysis to yield peaked likelihood profiles and relatively narrow CIs for all parameters, in particular supporting a sharp and significant past contraction signal with *N*_act/anc_ *<* 0.1.

**TABLE 1.**
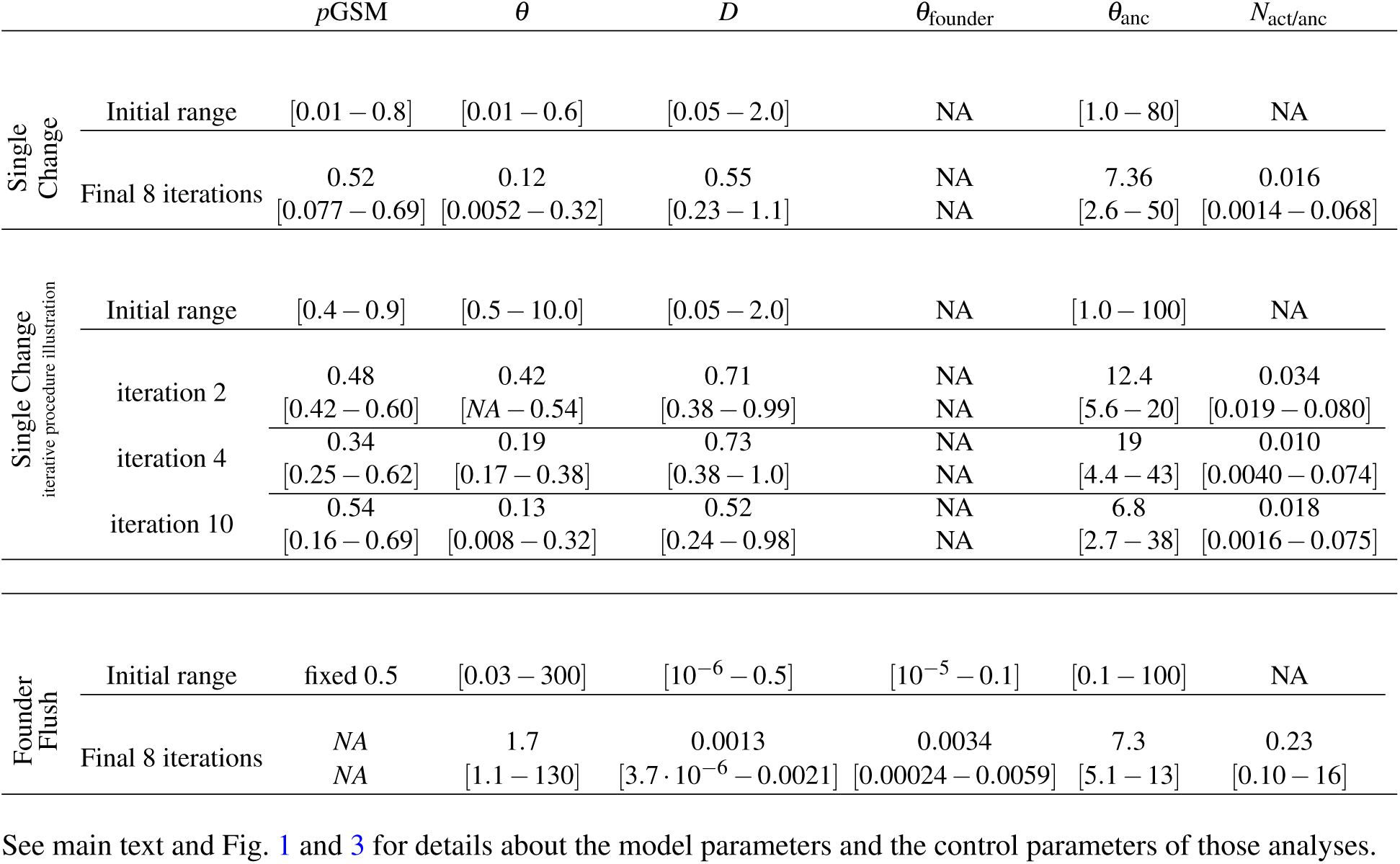
Initial parameter ranges, point estimates and 95% CIs obtained from the three analyses of the sheep data set.

**Figure 2:**
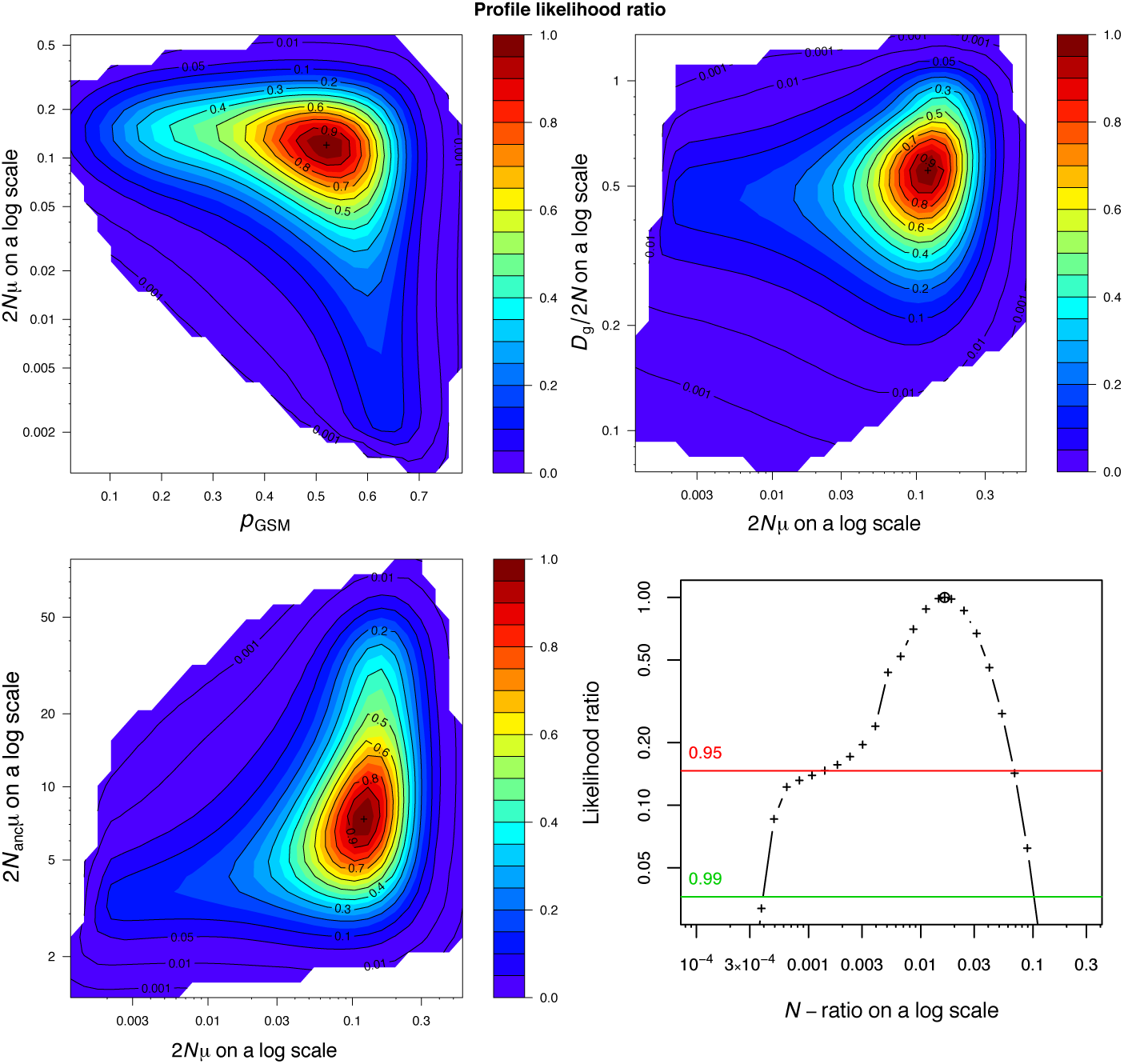
One- and two-dimensional profile LR plots for the sheep data set analyzed under the model with a single ancestral change in population size. See main text and Fig. 1 for details about the model parameters and the control parameters of the iterative analysis.

In a second step, we reanalyzed the sheep data under a more complex demographic model called “Founder-Flush” (FF), illustrated in Fig. 3. The FF model is designed for the analysis of samples from an isolated population that was founded some time in the past by an unknown number of individuals coming from a stable ancestral population of unknown size, and has then grown (or declined) exponentially until present (i.e., sampling time). Such a model is well suited to study invasive, reintroduced or epidemic populations and thus seems adapted to the sheep data set from Hirta.

**Figure 3:**
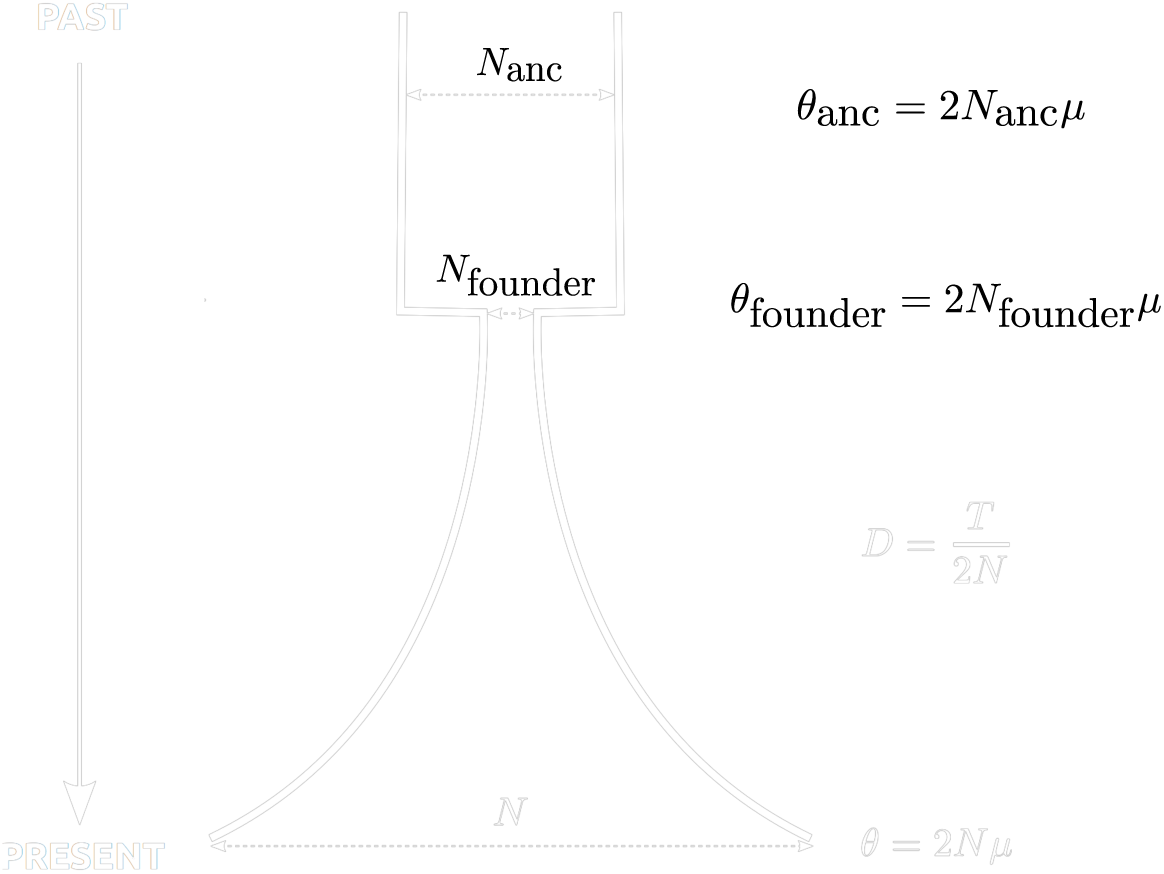
Representation of the Founder-Flush demographic model used for the analysis of the Soay sheep data set. *N* is the current population size, *N*_founder_ the size of the population during the founder event, and *N*_anc_ is the ancestral population size (before the demographic change), *T* is the time measured in generation since present and *µ* the mutation rate of the marker used. Those four parameters are the canonical parameters of the model. *θ*, *D*, *θ*_founder_ and *θ*_anc_ are the inferred scaled parameters of the coalescent approximation.

**Figure 4:**
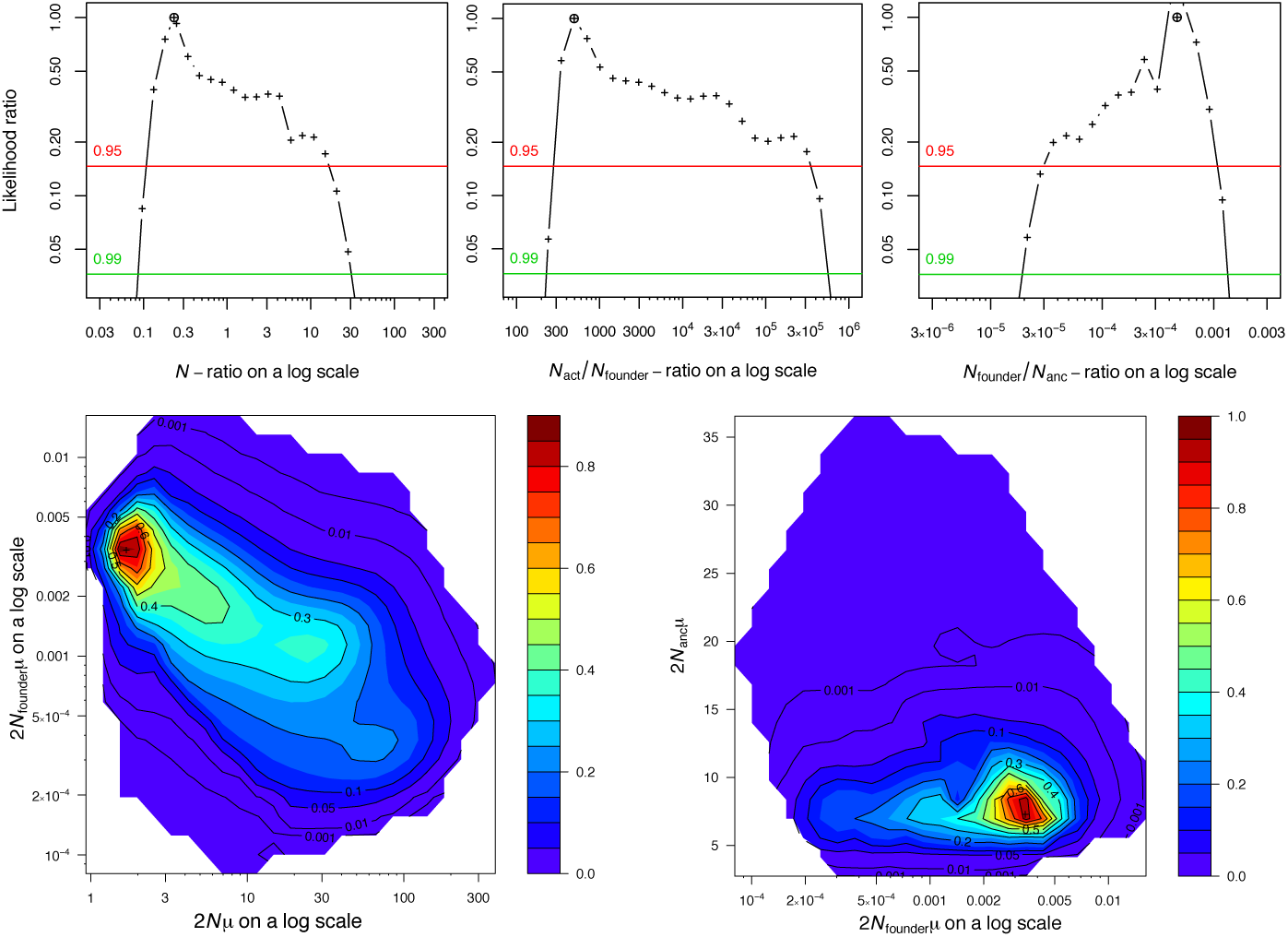
Examples of one- and two-dimensional profile LR plots for the sheep data set analyzed under the Founder-Flush model. See main text and Fig. 3 for details about the model parameters and the control parameters of the analysis.

As for the previous analysis, we considered a GSM model for mutations but we fixed its *p*_GSM_ value at 0.5 (i.e. the value inferred in the previous analysis) because preliminary analyses shows flatter profile likelihood surfaces when *p*_GSM_ is also estimated, thus complicating the whole analysis. This is probably due to the small number of loci (i.e. 17) of the data set, resulting in a lack of information about all parameters of the model. Four (scaled) parameters are thus inferred in this analysis: *θ* = 2*Nµ*, *D* = *T /*2*N*, *θ*_founder_ = 2*N*_founder_*µ*, and *θ*_anc_ = 2*N*_anc_*µ* (see legend of Fig. 3 for explanations of the model parameters). Three additional composite parameters are considered: (i) the *N*_act/anc_ = *N/N*_anc_ characterizing the ratio of the current population size vs the size of the source population; (ii) *N*_f/anc_ = *N*_founder_*/N*_anc_ characterizing the founder event; and (iii) *N*_act/f_ = *N/N*_founder_ characterizing the growth or decline of the newly founded population. The sheep data analysis under the Founder-Flush model was conducted by considering 8 iterations, with 300 points for which the likelihood is estimated using 2,000 genealogies. Initial parameter ranges as well as point estimates and associated CIs inferred after the 8 iterations are presented in Table. 1.

This second analysis under the FF model is coherent with the first analysis conducted under a simpler model: ancestral population size estimates are highly similar between the two analyses, with however narrower CI for the FF model. The FF analysis additionally detects the founding event (*N*_f/anc_ = 0.00047, CI: [3.4 · 10^*−*5^ 0.0010]). An expansion occurring after the founding event is also detected. Its estimate (*N*_act/f_ = 490, CI: [268 380, 000]) is higher than expected from census sizes, which may be due to difficulties in estimating population increases that are both large and recent, but other factors, such as variance in reproductive success, may also have strongly decreased the effective size of the founder population below its census size of 107 individuals. Finally, the inferred timing of the founder event is very recent (*D* = *T /*2*N <* 0.006), but coherent with the known time of introduction (i.e. 1932, corresponding to 19 sheep generations, given a generation time of four years as reported by Coulson et al., 2010, Table 3) and the inferred current population size. This is the first time to our knowledge that a founder-flush model, characterized by two ancestral changes in population size, is fitted using microsatellite loci. This analysis shows that small genetic data sets, as considered here, still contain relevant information about parameters of this model.

### 3.2. Adaptive exploration of likelihood surfaces

Inference of the likelihood surface uses an iterative procedure, as described in the previous section 2.2. Here, we illustrate this iterative procedure using the sheep data and the model with a single past change in population size as before, but considering bad initial ranges for two of the four parameters (*p*_GSM_ and *θ*). This shows the capacity of MIGRAINE to automatically adjust the sampled parameter space to regions of high likelihood that were not explored in the first iterations, and to gradually increase the density of points in those regions. For that purpose, the lower bounds of initial ranges for *p*_GSM_ and *θ* were both set at higher values than the corresponding CI lower bounds obtained in the previous analysis (see Table. 1).

Expectedly, the analysis with bad initial parameter ranges required more computation than the previous analysis to get satisfactory results. We doubled the number of points (i.e. 400) for which the likelihood is estimated at each iteration compared to the previous analysis and ran 10 iterations instead of 8. All other settings are identical. Table 1 presents point estimates and associated CIs for all inferred parameters at iterations 2, 4 and 10, and Fig. 5 represents the evolution of one-dimensional LR profiles through these iterations. Those results first show that, despite the bad initial parameter ranges, MIGRAINE succeeds in generating after 10 iterations point estimates and CIs similar to those obtained in the previous analysis with better initial parameter ranges. Additional iterations in both analyses only marginally change the results. Second, results from intermediate iterations show how MIGRAINE progressively extend the region of high likelihood. This automatic extension of explored parameter range is apparent in Fig. 6, which shows parameter points generated at iteration 2, whose likelihoods are to be estimated in the next iteration. Therein, different points generated according to different criteria are shown in different colors, and points generated by extrapolation in a previously unexplored parameter region are shown in black. The black points are mostly located at low *p*_GSM_ and *θ* values. The same points also have high *D* and *θ*_anc_ due to correlations between *p*_GSM_ and these two parameters near the likelihood maximum. This parameter correlation is apparent in Fig.6 and even more visible in the two-dimensional LR profiles for (*p*_GSM_,*D*) and (*p*_GSM_,*θ*_anc_) from the final iteration (results not shown). This diagnostic figure also shows that MIGRAINE samples points according to other criteria, aiming to ascertain the current likelihood maximum (orange points) and the CI bounds (red points), or to fill the region of high likelihood (roughly above the LR threshold of the confidence interval, but defined in two slightly different ways; green and dark blue points), or with high expected improvement outside this region (cyan points).

**Figure 5:**
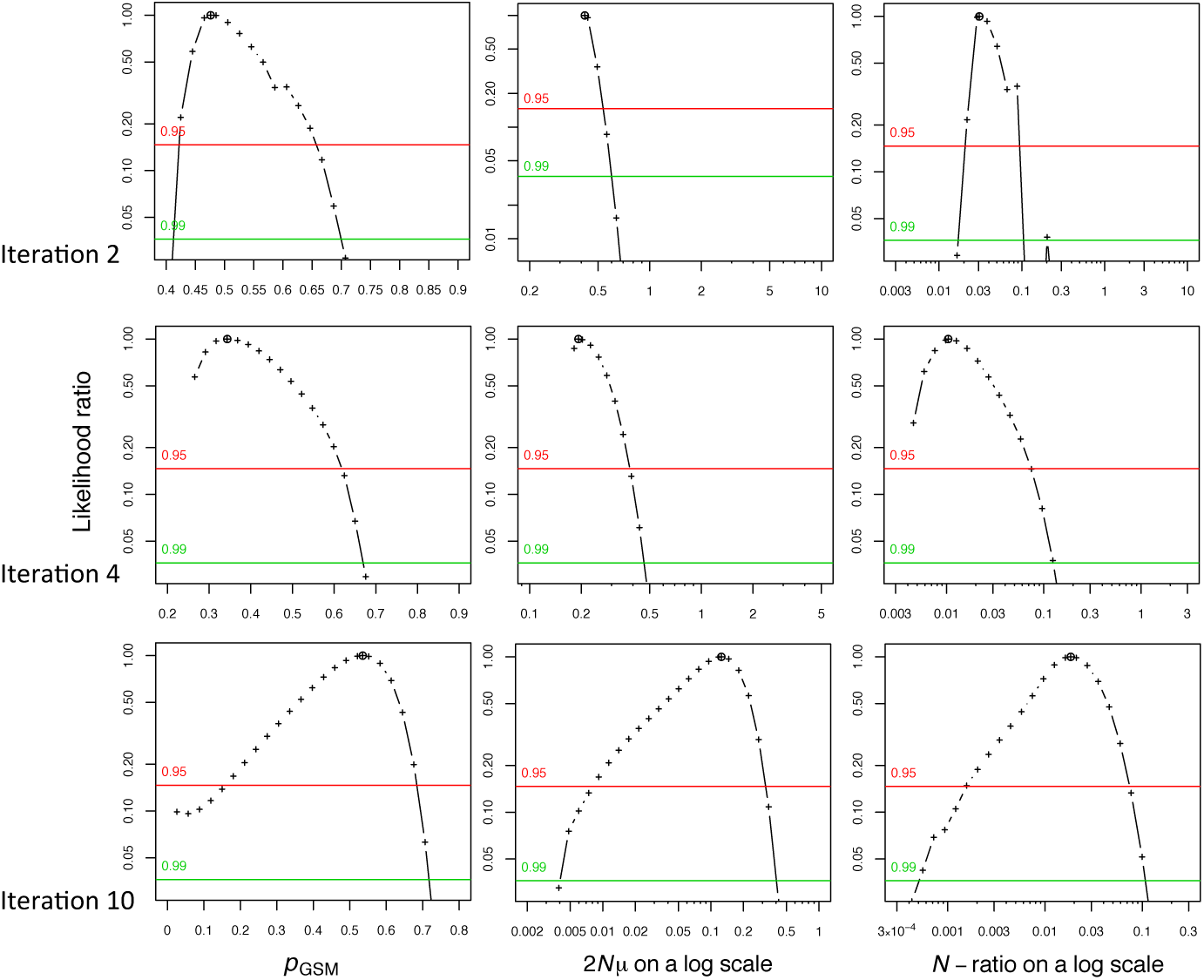
Illustration of the iterative procedure implemented in MIGRAINE for the sheep data set analyzed under the model with a single ancestral change in population size. Examples of one-dimensional profile LR plots for the parameters *p*_GSM_, *θ*, and *N*_act/anc_ for iterations 2, 4 and 10. See main text and Fig. 1 for details about the model parameters and the control parameters of the analysis.

**Figure 6:**
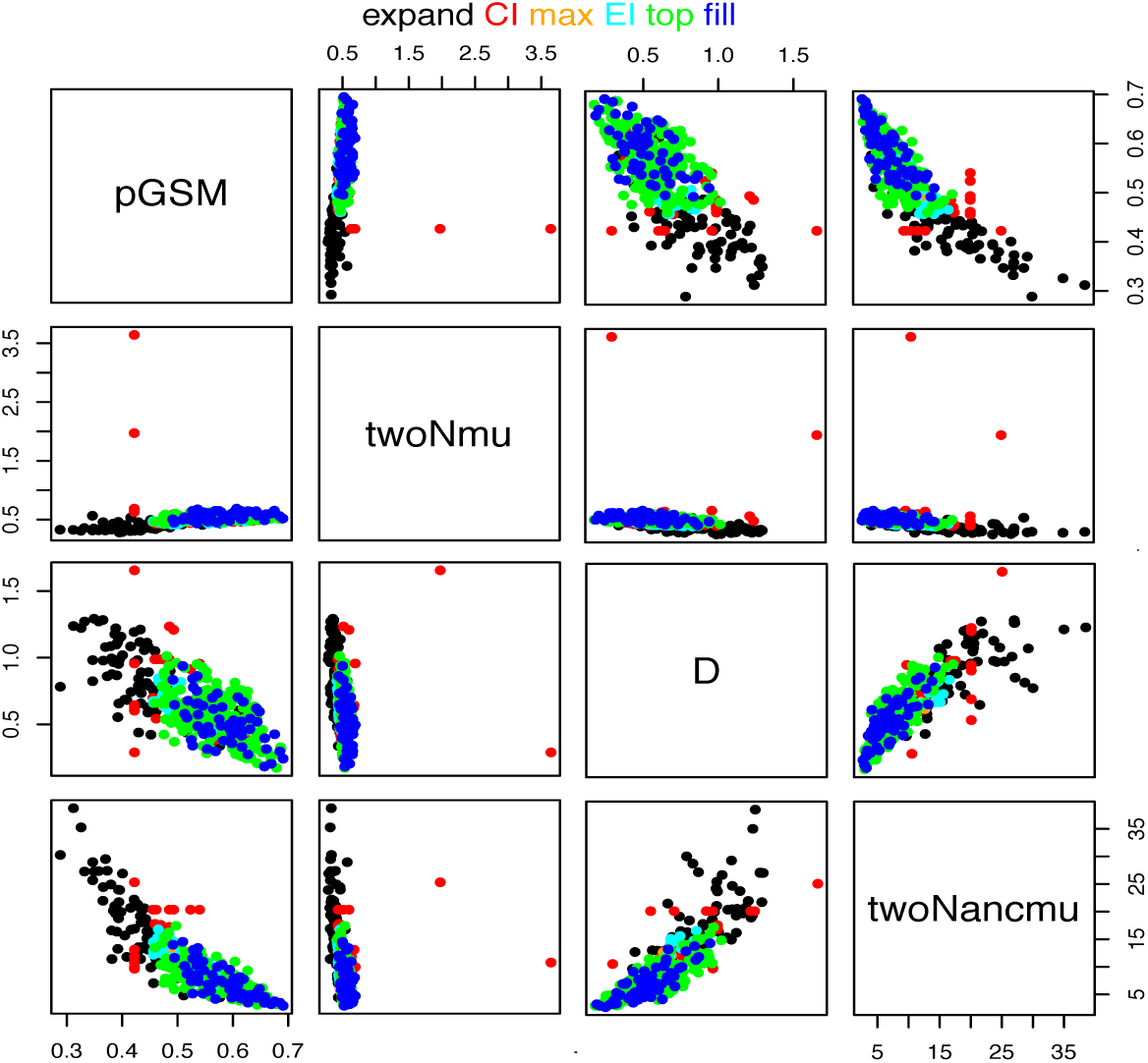
Diagnostic output graph illustrating the computation of new points during the iterative procedure implemented in Migraine. This graph shows the points defined at the end of iteration 2, for which the likelihood is to be estimated at iteration 3, for the sheep data set analyzed under the model with a single ancestral change in population size. See Main Text for the meaning of the different point colors. See main text and Fig. 1 for details about the model parameters and the control parameters of the analysis.

## 4. Discussion

### 4.1. Validation

The methods reviewed in this paper have been extensively assessed, in particular in terms of coverage of confidence intervals (e.g. Fig.7 and 8, and see Rousset and Leblois, 2012; Leblois et al., 2014). Assessment of coverage is not only suitable for interval estimation, but also more useful than assessment of bias and variance to detect problems in the inference of the likelihood surface by smoothing. Such assessment would hardly deserve mention, were it not for the fact that it is not the prevalent practice in broad segments of the literature related to this work either in its objectives (inference from genetic variation) or through its methods (various stochastic methods to infer likelihoods or posterior distributions). Consequently, poorly assessed methods or software are readily available and endorsed by practitioners eager to make a story out of their data. In fact, very few publications testing methods for population genetic inference even mention confidence intervals coverage properties. Moreover, the few papers that report such information often find strong inaccuracies of the CIs (e.g. Abdo et al., 2004; Beerli, 2006; Hey, 2010; Hey et al., 2015; Appendix S3 of Peter et al., 2010).

The method defined by de Iorio and Griffiths (2004a,b) provides an approximation for the probability that a newly sampled gene is of a given type. As noted above, this approximation reduces, under a model of a single stationary population with parent-independent mutations (PIM, i.e. when the forward mutation rate from genetic type *i* to *j* is independent of *i*), to the true probability, and thus leads to the optimal importance sampling distribution, allowing “perfect simulation” under such a model. Under stationary models of structured populations, this approximation does not allow perfect but still very efficient importance sampling simulation.

Imperfect performance of the inferences can still result from approximations inherent in the methods. In our own work, examples include biased estimation of parameters when the analytical approximations inherent to coalescent and diffusion approximations (e.g., large population size) do not hold (Rousset and Leblois, 2012), poor robustness of some inferences with respect to details of the spatial organization of the population (Rousset and Leblois, 2007, 2012), and large variance of the importance sampling algorithm in non-equilibrium models (Leblois et al., 2014).

Poor performance could also be expected because the traditional assumptions of asymptotic likelihood theory do not hold. A first reason is the discrete nature of the data, which occasionally impacts the distribution of the likelihood ratio even in large samples. To understand how this can occur, first consider the infinite allele model (IAM), according to which each mutation generates an allele not preexisting in the population. In this model, the observed number of alleles *k* in a sample is a sufficient statistic for *θ* (Ewens, 1972), and as it is a discrete variable, the distribution of the LR is also discrete. The IAM may be seen as a limit case of the *K*-allele model (KAM), a model with *K* possible allele types and identical mutation rates between any pair of alleles.

For the KAM with large *K* (thus approaching the IAM), but with a small mutation rate (thus with few likely values of *k*), steps in the distribution of the likelihood may thus become visible. This is illustrated in Fig. 7, which shows the analysis of samples of 100 gene copies generated under a 20-allele KAM. Steps are visible when these data are analyzed under a KAM with large K (400; Fig. 7a), while they disappear for small *K* (20; Fig. 7b). A second and more general deviation from traditional assumptions is that a sample of *n* genes is typically not considered as resulting from *n* iid draws. Instead, the *n* genes are related through their common ancestry, and the realized ancestral genealogy can be viewed as a single draw of a latent variable. The impact of this dependence is clear for example on the inference of the mutation rate under the infinite allele model (IAM), where the variance of the ML estimator is asymptotically *O*[1*/* log(*n*)] rather than *O*(1*/n*) (Tavaré, 1984, p. 41). Yet, in the KAM model with small *K*, we can achieve practically perfect coverage from small samples (*n* = 30 genes from a single locus, Fig. 7b). This observation, coupled with the fact that it is recommended to apply such methods to samples of several unlinked loci, suggests that the genealogical dependence has little impact on likelihood approximations.

**Figure 7:**
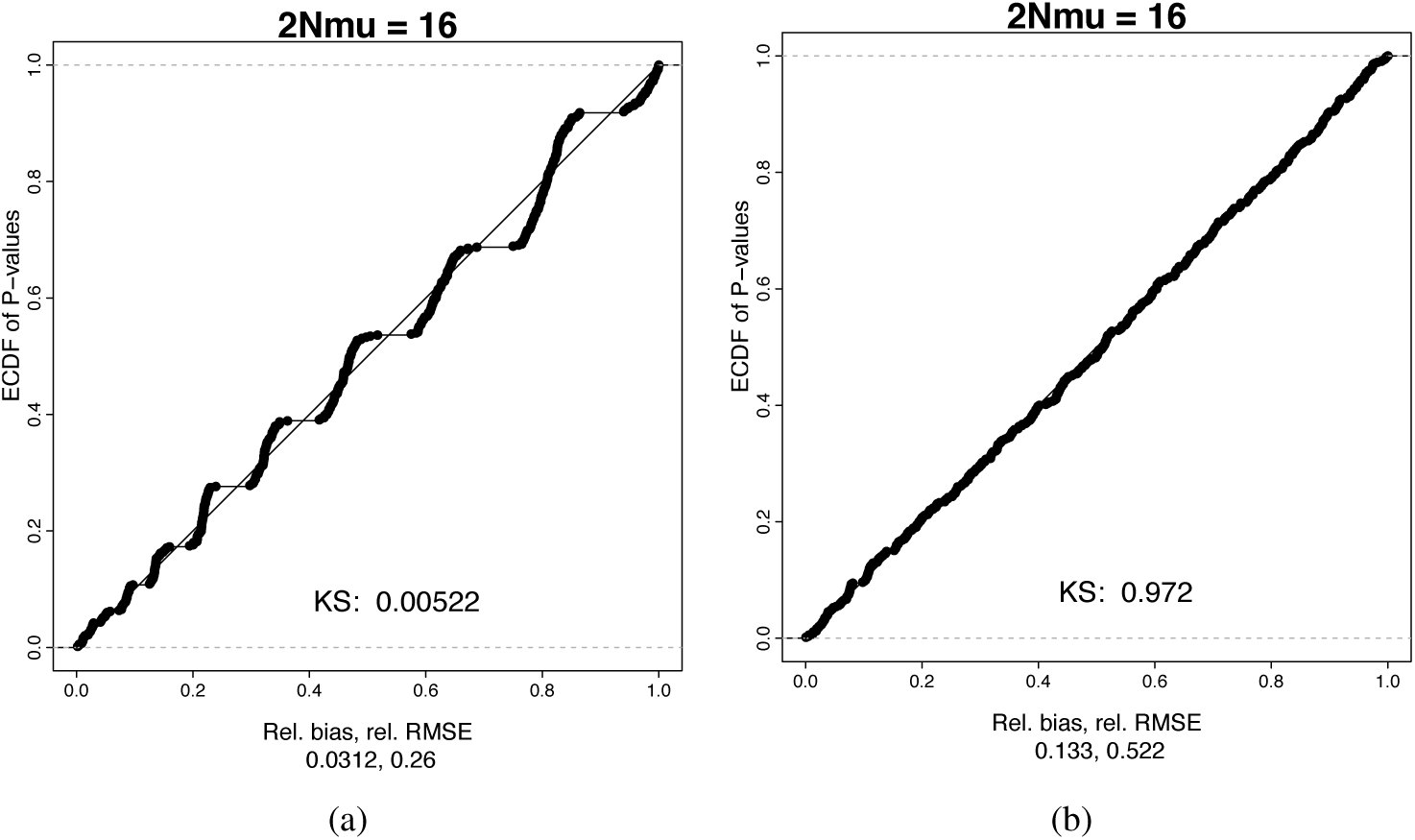
Empirical cumulative distribution functions (ECDF) of P values of LR tests under a model of a single stable population *θ* = 2*Nµ* = 16.0 for a KAM with (a) *K* = 400 and (b) *K* = 20 possible alleles. Mean relative bias (rel. bias, computed as 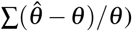 and relative root mean square error (rel. RMSE, computed as 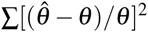 are reported. KS indicate the P value of the Kolmogorov-Smirnov test for departure of LRT P values distributions from uniformity.

### 4.2. Drivers of robustness under imperfectly specified models

None of the mutation models implemented can be considered as exact representations of the actual mutation processes at the markers assayed. Thus, one typically considers a simple model such as the PIM in order to make inferences about other parameters such as dispersal rates. Robustness has been checked in this case (Rousset and Leblois, 2012). Isolation by distance analyses under a PIM model of microsatellite data simulated under a strict stepwise mutation model (SMM, Ohta and Kimura, 1973), according to which mutation results in the gain or loss of only one repeat of the DNA motif, showed that mis-specification of the mutation model has little impact on dispersal estimator performance, but a 50 to 75% bias in scaled mutation rate estimates is observed (Rousset and Leblois, 2007, 2012). This bias is expected because the variation in local diversity in KAM versus SMM is approximately that resulting from a 2-fold variation in mutation rate (Rousset, 1996). Similarly, inference of scaled migration rates between pairs of populations, but not of scaled mutation rate, is expected to be robust to mutational processes.

On the other hand, inferences in demographic models with time-varying parameters are much more sensitive to mutational processes. Leblois et al. (2014) showed that mis-specification of microsatellite mutational processes can induce false detection of past contraction in population sizes from samples taken from stationary populations. It can also induce biases in inferred timing and strength of a past change in population size from samples taken from a population that has indeed undergone past demographic changes. We have thus implemented variants of the importance sampling algorithms for different mutation models. Such work is illustrated for an unbounded SMM and models with one or two populations, in de Iorio et al. (2005), in Leblois et al. (2014) for a generalized stepwise mutation model (GSM) in a single population, and in Fig. 8 for the Infinitely many Site model (ISM; Kimura, 1969), a model adapted to DNA sequence markers (see next section).

**Figure 8:**
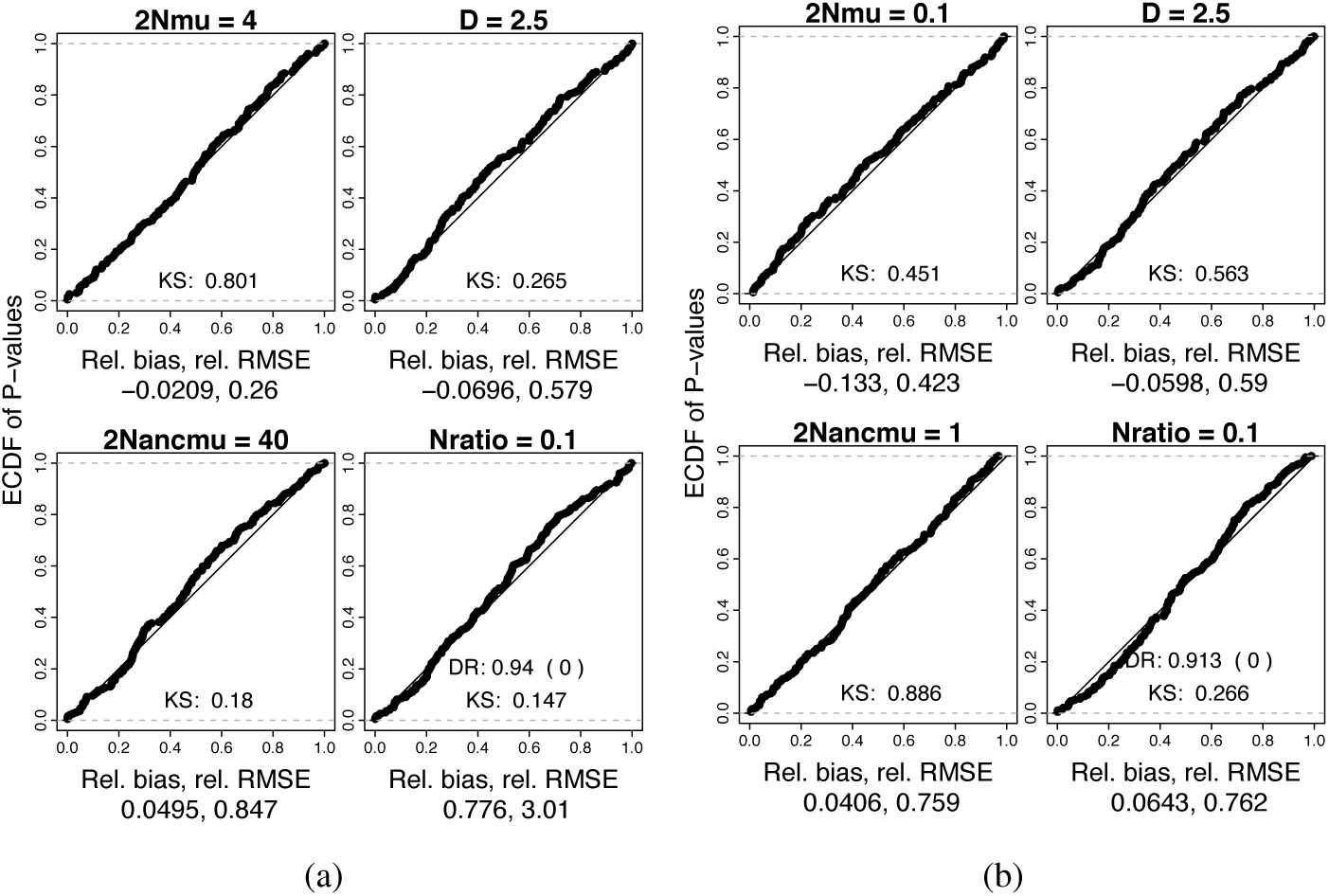
ECDF of P values of LR tests for a scenario with a single past change in population size as illustrated in Fig. 1 (a) for 30 SMM loci with *θ* = 2*Nµ* = 4.0 and and *θ*_anc_ = 2*N_anc_µ* = 40.0; (b) for 30 ISM loci with *θ* = 2*Nµ* = 0.1 and and *θ*_anc_ = 2*N_anc_µ* = 1.0; *D* = *T /*2*N* = 2.5 for both analyses. Mean relative bias (rel. ^bias, computed as^ ∑(observed value parameter value)*/*parameter value) and relative root mean square error (rel. RMSE, computed as ∑[(observed value parameter value)*/*parameter value]2 are reported as well as the contraction detection rate (DR) and false expansion detection rate (FEDR) in parentheses after DR. KS indicate the P value of the Kolmogorov-Smirnov test for departure of LRT P values distributions from uniformity.

### 4.3. Expected developments

All the mutation models discussed above describe allelic data, typically microsatellites, and not DNA sequences, which are also widely used genetic markers. Even if non-recombining DNA sequences can be analyzed as allelic data by considering haplotype identity only, it implies a great loss of genetic information carried by the mutations present in the different haplotypes. One mutation model adapted to DNA sequences, the infinitely-many-site model (ISM, Kimura, 1969), has been considered since the earliest developments of coalescent-based importance sampling algorithms, e.g. in the software GeneTree developed by Bahlo and Griffiths (2000) and in the approximations defined in de Iorio and Griffiths (2004b). Hobolth et al. (2008) also developed a specific proposal distribution based on an exact sampling formula from a single DNA site. Simulations showed however that the latter proposal is not more efficient than de Iorio & Griffiths’ ISM specific solution derived from their general approximation (unpublished results). Nevertheless, both proposals for the ISM model have been implemented in MIGRAINE and have already allowed analysis of real data sets with sequence data (e.g., Vignaud et al., 2014b, Lalis et al., 2016). Extensive simulation tests of the ISM implementation in MIGRAINE are not yet published, but we show in Fig. 8b good performances, in terms of relative bias, relative RMSE and coverage properties of the CIs, of such analyses of DNA sequence markers evolving under the ISM compared to microsatellite markers evolving under the SMM (Fig. 8a) under a scenario with a single past change in population size (i.e., the model presented in Fig. 1).

Finally, given the explosion of single nucleotide polymorphism (SNP) data, it would be interesting to develop IS algorithms specifically adapted to SNPs, but except for de Iorio and Griffiths’ suggestion to use the ISM algorithm with a single site and let *θ* parameters tends to 0, we are not aware of any development, application or test of IS algorithms for SNP data. SNP data may also be analyzed under a KAM with two possible alleles for the inference of dispersal between subpopulations because such inference is robust to mis-specifications of the mutation processes. On the contrary, inferences under time-inhomogeneous models may be strongly biased by such model mis-specification, especially for the timing of the different ancestral events (e.g. changes in population or divergence events).

But all algorithms dedicated to the alternative mutation models increase computation time of each replicate in comparison to the PIM, and for a given number of replicates, none has exhibited a variance as low as algorithms defined for the PIM. The current approaches for designing importance sampling algorithms are less and less efficient when mutation is more dependent on the parental type: as reviewed above, they work best for the PIM, then the GSM and the SMM, and the ISM comes last here.

### 4.4. Conclusion

The works reviewed here have shown the feasibility of likelihood-based inference for an increasing range of models of data types and demographic processes. A broader range of inferences (e.g., in demographic models with large rates of coalescence or migration) may be currently prevented by the limitations inherent to the approximations of coalescent and diffusion approaches. Analyzing a large number of loci (e.g. few thousands for typical NGS data on non-model organism) may also be challenging because of (i) the additive effect of the variance observed at each locus; and (ii) potentially large computation times. Such limitations underlie the persistent scope for alternative methodologies. Even within the current framework, there is still scope for substantial improvements, in particular of importance sampling algorithms for specific mutation models and time-inhomogeneous demographic models.

## Acknowledgements

We thank Jean-Michel Marin for inviting this contribution, and Josephine Pemberton for sharing her data from the sheep population from Hirta island. Part of this work was carried out by using the resources of the INRA MIGALE (http://migale.jouy.inra.fr) and GENOTOUL (Toulouse Midi-Pyrénées) bioinformatics platforms, the computing grid of the CBGP lab and the Montpellier Bioinformatics Biodiversity platform services. This study was supported by the Agence Nationale de la Recherche (project IM-Model CORAL.FISH 2010-BLAN-1726-01) and by the Institut National de Recherche en Agronomie (Project INRA Starting Group “IGGiPop”, and postdoctoral funding for C. R. Beeravolu).

## 5. Appendix

An efficient importance sampling algorithm has been formulated using concepts from diffusion theory. We first recall how diffusion models are used in population genetics (see Ewens, 2004 for an extensive introduction).

A process of allele frequency change in a finite population of size *N* with (say) mutation rate *µ* is approximated by the limiting process as *N* → ∞, of a series of processes *X*_*N*_ with the same *Nµ*, each measured in time scaled by *N*. For example, consider a locus with only two possible alleles, with mutation probability *µ* to the other allele per gene copy per generation. The expected allele frequency in the next generation is E(*x^′^*) = *x*(1 −*µ*) + (1 −*x*)*µ*. in the classical Wright-Fisher model for a population of *N* haploid individuals, the allele frequency in the next generation is a binomial sample of size *N* with binomial frequency E(*x*′) as given above. The change in allele frequency has then expectation E(*x*′) *x* = (1 −2*x*)*µ* and variance E(*x*′)(1 −E(*x*′))*/N*. The limiting process in scaled time is then described by the infinitesimal moments in scaled time in units of *N* generations, *M*(*x*) = *Nµ*(1 − 2*x*) and *V* (*x*) = *x*(1 − *x*). In particular, the transition density *φ* (*X_t_ X*_0_) of allele frequency in the limiting process satisfies the backward Kolmogorov equation

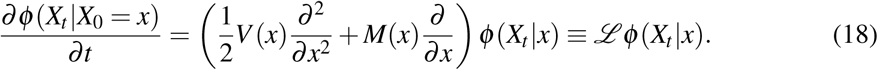

A backward equation holds also for the expectation of any function *f* (*x*) with bounded second derivatives (“generator equation”; Karlin and Taylor, 1981, p. 215),

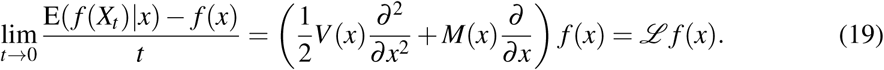

These results are extended to models with multiple alleles and subpopulations, with the following notations. We consider deme sizes, *N*_*d*_ for deme *d*, which sum to *N*_T_; a matrix of scaled forward mutation rates *N*T*µi j ≡ N*T*µPi j* from *i* to *j* (which is row-stochastic, i.e., ∑ _*j*_ ^Pi j^ = 1); and a matrix of scaled forward migration rates *N*_T_*m*_*dd*_^*−*^ from deme *d*′ to *d*. the diffusion process is now the limit, as *N* → ∞, of a series of processes *X*_*N*_ with the constant *Nµ*_*i*__*j*_, constant *Nm*_*dd*_′, and constant relative deme sizes. Then the generator can be written

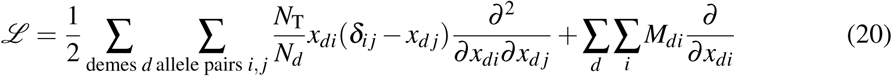

where 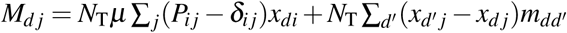

At stationary equilibrium, 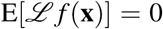 where **x** ≡ (*x*_*id*_) is the vector of frequencies of allele *i* in deme *d*, and expectation is taken over the joint stationary density *ψ*(**x**) of these allele frequencies. Applying this result for *f* taken as the sample probability given **x**, i.e. 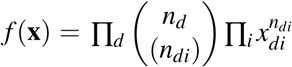 where *xdi* is the frequency of allele *i* in deme *j*, leads to a relation between probabilities of samples that differ by one coalescence/mutation/migration event:

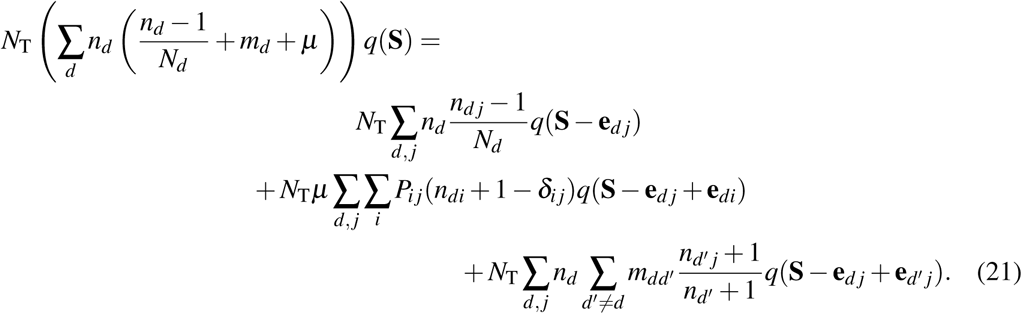

We use eq. 13 to express eq. 21 as a recursion involving ancestral samples differing by the subtraction of one gene copy relative to the descendant sample, by expressing all *q*(.) in terms of *q*(**S** *−* **e**_*dj*_)s for distinct *d, j*:

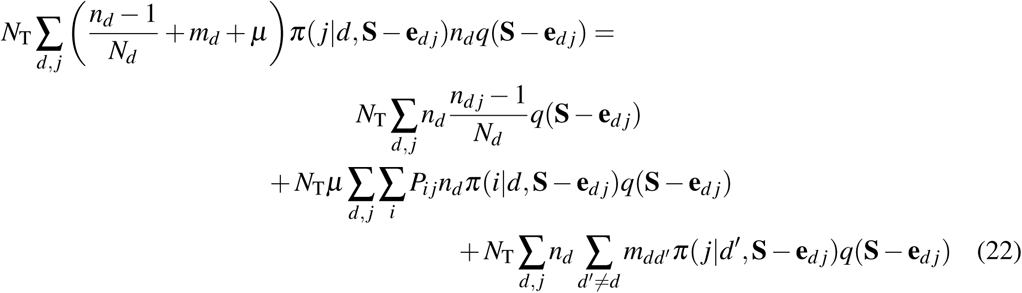

This provides no solution for the *π*’s, as the closed system of equations for sample probabilities implied by this one is not simplified in any way, and remains too large. But de Iorio and Griffiths (2004b) instead considers the equations defined for each *d, j* by extracting the left-hand and right-hand side coefficients of *q*(**S − e**_*dj*_). The system of such equations for different samples of same size as **S** over all *d* and *j* is generally inconsistent. However, the system of equations for identical *q*(**S − e**_*dj*_) over different *d, j* leads to a linear system of equations of dimension the number of demes times the number of alleles. Each such equation reduces to

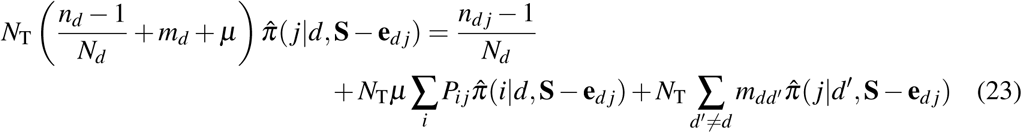

where e.g. 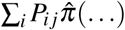 represents a sum over different possible ancestral sample configurations with an additional *i* gene, cf eq. (21). The 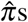 solving this system are not the true *π*s, but they provides approximations for the *π*s from which importance sampling weights and a proposal distribution can be defined.

